# The locus coeruleus directs sensory-motor reflex amplitude across environmental contexts

**DOI:** 10.1101/2022.05.25.493447

**Authors:** Emily C. Witts, Miranda A. Mathews, Andrew J. Murray

**Author notes:** Corresponding authors, (AJM) or (EW).

## Abstract

Animals possess a remarkable ability to quickly and accurately respond to challenges to their balance and posture. Postural corrections are the implementation of a motor act by the nervous system that counteracts a perturbation and returns the body to a stable state. These corrections must respect both the current position of the limbs and trunk, as well as the external environment. However, how motor circuits integrate multiple streams of information regarding both these internal and external factors, and adjust motor actions accordingly, are poorly understood. Here we show that the lateral vestibular nucleus in the brainstem generates motor corrections following perturbation, and that this reflex can be altered by manipulating the surrounding environment. The strength of the motor correction is influenced by noradrenergic signalling from the locus coeruleus, suggesting a potential link between forebrain structures which convey sensory information about the environment, and brainstem circuits that generate motor corrections.

## Introduction

To navigate unpredictable environments animals and humans must be able to quickly and accurately respond to disruptions in balance and posture. Upon sensing an external perturbation, such as a shift in support surface or mechanical impact, the nervous system must promptly interrupt ongoing motor plans and implement an action that maintains upright posture. The goal of this action is to precisely counteract the perturbation and maintain the centre of mass (CoM) within the base of support (Carpenter et al 1999a, Deliagina et al 2008, Denny-Brown 1929). This response must respect both the current position and limitations of the body, as well as the surrounding environment (Balasubramaniam & Wing 2002, Cleworth et al 2012, Macpherson et al 1989). This seemingly simple action is thought to be implemented by a combination of segmental reflexes in the spinal cord, along with vestibulospinal and reticulospinal pathways in the brainstem and can be influenced by forebrain structures such as motor cortex (Deliagina et al 2007, Jacobs & Horak 2007, MacKinnon 2018, Murray et al 2018).

Postural reflexes, however, are not simple transformations of the sensory input signalling an unexpected movement into an opposing motor response. The motor plan generated in response to a perturbation can be heavily influenced by the animal’s surrounding environment, even when that environment has no mechanical influence on the perturbation itself. Intuitively, we know that the presence of a nearby handhold will have a different influence on our motor response than if there were a steep drop on that same side. This environmental effect has been considered as a cortical influence on the sensorimotor set, which can prime motor responses to a perturbation (Beloozerova et al 2005, Jacobs & Horak 2007). In a similar, but simpler, scenario, postural responses can be influenced by factors that alter threat levels in humans. When human subjects stand at the edge of a high surface, they alter their response to perturbations of the support surface (Carpenter et al 2001) and adopt a stiffening strategy that limits movement and postural sway (Carpenter et al 1999b). Functionally, this behavioural response may act to limit the chances of severe injury that would result from falling from height (Horslen et al 2018). At a neural circuit level, postural threat has been linked with altered sensitivity of the muscle spindle (Horslen et al 2013), altered central processing of vestibular input (Horslen et al 2014) and a change in gain in vestibulospinal reflexes (Naranjo et al 2015, Naranjo et al 2016). These studies are generally in agreement with work done in the mouse showing that optogenetic stimulation of the lateral vestibular nucleus (LVN) only results in motor responses when the animal is on a balance beam at height, and not when walking on the stable surface of a treadmill (Murray et al 2018). In general, this ability to flexibly modify postural responses across terrains and environmental conditions is a critically important component of the balance system and a lack of flexibility of this system may contribute to destabilisation in disorders such as Parkinson’s disease (Horak et al 1992, Rinalduzzi et al 2015).

The question remains as to how neural circuits that encode environmental information, such as threat level, influence the brainstem and spinal circuits that generate postural reflexes. Environmental influences on postural control have predominantly been studied in humans, thus providing a relatively limited understanding of these pathways. To address this, we developed a novel behavioural paradigm which allowed us to alter the surrounding environment of a mouse undergoing a postural perturbation without altering the mechanical aspect of the perturbation itself. Here, kinetically identical perturbations resulted in differing motor responses at both the behavioural and muscle level. Using a combination of behavioural analysis, pharmacology and selective circuit manipulations we found that noradrenergic signalling from the locus coeruleus influences the output level of LVN-dependent postural corrections and may therefore provide an important link between cognitive circuits that interpret environmental context and the motor pathways that maintain balance.

## Results

### Environment influences the motor response to a perturbation

Vestibulospinal reflexes act to maintain balance and upright posture. The LVN, and its spinal projection from lateral vestibulospinal tract (LVST)-neurons, is a key brainstem structure required for the generation of a reflexive motor correction following a postural perturbation (Murray et al 2018). To ascertain whether these postural reflexes could be influenced by environmental context we developed a behavioural paradigm that could alter perceived threat levels as mice underwent a perturbation.

Rodents are known to avoid open spaces due to the possibility of predation and show a preference for enclosed spaces, particularly in novel environments (Ennaceur 2011). This is exemplified in rodent behavioural tests of anxiety such as the elevated plus maze or thigmotaxis in an open field arena (Crawley & Goodwin 1980, Graeff et al 1998, Kulesskaya & Voikar 2014). Using this information as a starting point we designed a behavioural paradigm where mice would undergo a lateral postural perturbation forcing them to make a corrective motor response to maintain an upright posture. Mice received this perturbation in an enclosure measuring 10 cm by 6 cm in the XY plane. In order to test whether the external environment could influence the postural response we varied the height of the enclosure in the Z plane. A “low walled” condition where the height of the enclosure was slightly above head height of the animal was compared to a “high walled” condition which was double this height (3.5 vs. 7 cm; Figure 1A). We hypothesised that mice would feel more exposed or threatened in the low walled condition. Consistent with this, we observed the respiratory rate of mice (a proxy for anxiety; (Adhikari 2014) to be elevated in the low wall condition (220.5+/-7.5 breaths per minute) compared to high walled condition (180.1+/-14.6 breaths per minute; p=0.049; Figure 1B). A perturbation was then introduced by rapid lateral movement of the entire arena. We measured the movement of each arena to ensure animals received identical perturbations in the two environments. The displacement of each arena was 115 mm in the lateral plane with a peak acceleration between 0.53 and 0.54 ms, reached 50 ms from movement onset. The entire movement lasted 140 ms (Figure 1C; Supplementary Figure 1A,B).

**Figure 1:**
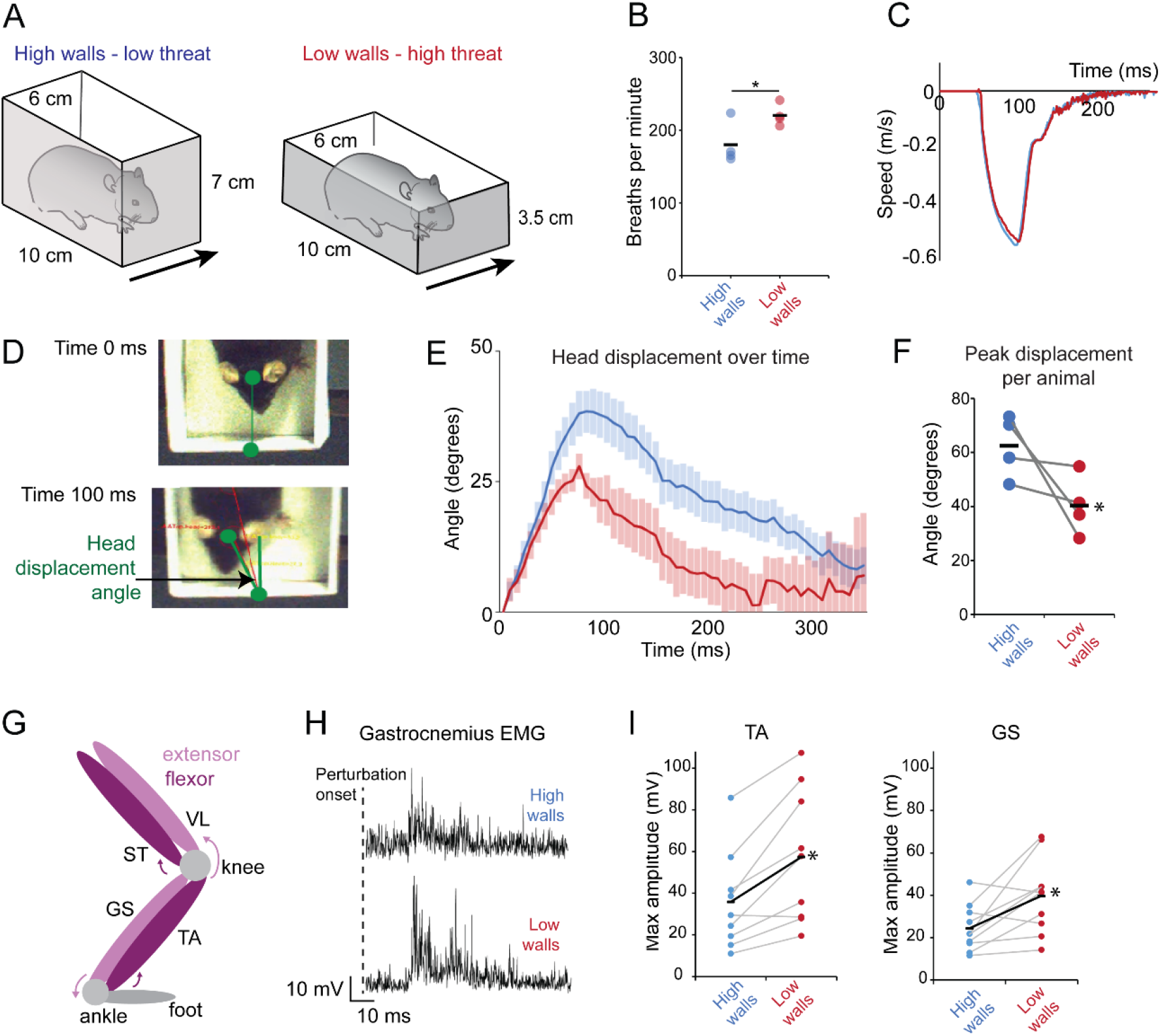
Environmental context alters response to unexpected perturbation. (A) Schematic showing the behavioural apparatus to vary environmental context. (B) Breaths per minute in experimental animals in the high and low wall conditions. (C) Traces of the arena displacement (perturbation) in the high and low walled conditions. (D) Example images of head position relative to fixed point in the arena prior to perturbation onset (Time 0) and 100 ms after perturbation onset. (E) Change in head displacement angle over time from perturbation onset (time 0) in high and low walled conditions. (F) Peak head displacement angle after perturbation in high and low walled conditions. (G) Diagram showing location of hindlimb muscles implanted with electrodes (see also supplementary figure 1) (H) Example rectified EMG traces from the GS muscle in high and low walled condition after perturbation onset. (I) Peak EMG amplitude of TA and GS muscles during perturbation in high and low walled conditions. Each point is the mean of trials from individual experimental animals with the overall mean represented by black lines. *p<0.05.

We closely monitored the postural correction initiated by the mice following the perturbation. We measured the displacement of a fixed point on the head over time starting at perturbation onset (time 0; Figure 1D,E; Supplementary Figure 1B), relative to a point within the arena. Animals in the low walled environment were displaced significantly less than those in the high walled condition (Figure 1E) consistent with a stronger, and more effective, postural correction. The peak displacement of the head (defined as the maximum angle reached by the head compared to a fixed point on the arena) for the high walled condition was 62.4+/-5.8° compared with only 40.3+/-5.5° for the low walled condition (p=0.032; Figure 1F). Similarly, fixed points on the body and base of the tail showed decreased stability in the high walled condition over the 400 ms recording period (Supplementary Figure 1C, D).

To examine limb muscle activity underlying the difference in body movement, we performed EMG recordings from the extensor muscles gastrocnemius (GS) and vastus lateralis (VL) and flexor muscles tibialis anterior (TA) and semitendinosus (ST) muscles in the hindlimb (Figure 1G). Consistent with a stronger postural correction, the peak EMG amplitude in ankle muscles GS and TA were larger in the low wall condition (39.7 ± 5.6 mV; n=10 mice and 57.4 ± 10.7 mV; n=9 mice) compared to the high wall condition (24.4±3.4 mV; n=10 mice; p=0.0023 and 35.9±7.9 mV; n=9 mice; p=0.016; Figure 1H,I; Supplementary Figure 1F). The latency of onset of the EMG response (measured as time from perturbation onset to the peak response; Supplementary Figure 1E) was not different between conditions, indicating that only the amplitude and not the timing of the EMG response was influenced by the environment.

### Silencing of the lateral vestibular nucleus prevents postural corrective responses

Vestibulospinal pathways convey proprioceptive and vestibular sensory information to spinal motor circuits (Witts & Murray 2019) and these pathways are required for postural corrections in mice (Murray et al 2018). However, local reflexes at the level of the spinal cord also contribute to postural control (Deliagina et al 2008). To test whether the LVN was required for the generation of postural reflexes in our behavioural assay we inhibited the activity of the LVN during lateral perturbations (Figure 2A-D). We stereotaxically injected an adeno-associated virus (AAV) expressing the outward proton pump archaerhodopsin (ArchT;(Chow et al 2010)) into the LVN and implanted a fibre optic probe to deliver yellow light to neurons in this area (Figure 2A,B), selectively and reversibly inhibiting the LVN during postural perturbations.

**Figure 2:**
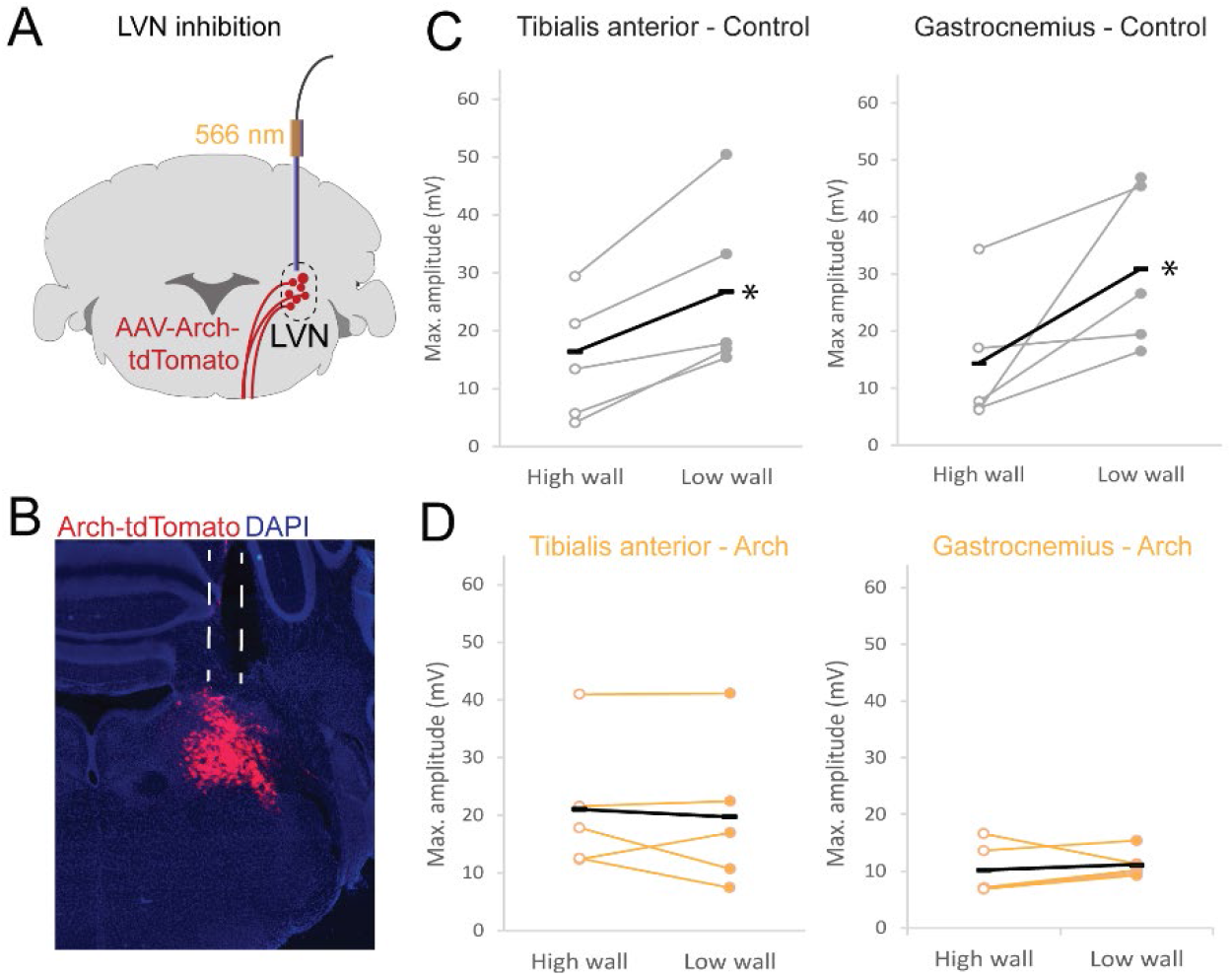
Inhibition of the LVN reduces response to perturbation. (A) Schematic showing experimental strategy to transiently inhibit neurons in the LVN. (B) Histological image showing virus injection in the LVN and placement of fiber optic cannula. (C) Mean peak EMG amplitude after perturbation without light (control). (D) Mean peak EMG amplitude after perturbation with light on and inhibition of LVN neurons. In B and D each point is the mean of trials from individual experimental animals with the overall mean represented by black lines. LVN-lateral vestibular nucleus. *p<0.05.

When neurons in the LVN were inhibited using archaerhodopsin, the response to a lateral perturbation was substantially altered. In TA, without light, there was an elevated response in low wall (26.8±6.7 mV; n=5 mice) compared to high wall (14.8±4.7 mV; n=5 mice; Figure 2C) conditions, as observed above. However, in the same animals, the presence of light to silence cells in the LVN caused a reduced response to perturbation in both the high wall (21.4±6.3 mV; n=5 mice) and low wall (19.7±5.9 mV; n=5 mice) conditions (Figure 2D). Similarly, in GS, in the absence of light, amplitude of EMG signals in response to perturbation was larger in the low wall condition (30.9±6.4 mV; n=5 mice) compared to the high wall condition (14.3±5.4mV; n=5 mice; Figure 2C). When the light was on, however, and neurons in the LVN were inhibited, there was a reduced response in both the low wall (11.1±1.1 mV; n=5 mice) and high wall (10.2±2.0 mV; n=5 mice) conditions (Figure 2D). The extent of inhibition in ArchT-expressing neurons can be altered by varying the light power (Chow et al 2010). Increasing the light intensity whilst recording EMG responses to perturbations resulted in a reduced EMG response in both high and wall conditions, consistent with increasing inhibition of motor signal from the LVN (Supplementary Figure 2).

### Noradrenergic signalling contributes to normal locomotor activity

The results above, along with previous work (Murray et al 2018, Pompeiano 2001), indicate that vestibulospinal neurons are required for the muscular response to counteract a perturbation. Our behavioural paradigm indicates that this response can be varied dependent on environmental context, but how does information regarding the surrounding environment feed into postural circuits?

One possibility is the involvement of noradrenergic signalling. Previous studies looking at vestibulospinal reflexes in decerebrate cats have indicated that noradrenaline can influence vestibulospinal reflex gain (Fung et al 1991, Pompeiano 1989, Pompeiano 2001). The locus coeruleus (LC) has been suggested to be the source of this noradrenergic input to the LVN and is also known to have further roles in overall posture and arousal (Carter et al 2010). Given that the LC is implicated in attention and vigilance (Sara 2009), we hypothesised that this region could provide the necessary link between environmental context and postural control pathways.

Studies in reduced preparations have demonstrated that noradrenergic activity is involved in the initiation and modulation of locomotion (Miles & Sillar 2011) and can directly influence motor neuron activity (Hultborn & Kiehn 1992). Therefore, prior to assessing postural control, we first examined whether blockade of noradrenergic signalling resulted in any gross motor changes. To achieve this we selectively disrupted noradrenergic signalling with the compound N-(2-chloroethyl)-N-ethyl-2-bromobenzylamine (DSP-4; (Jaim-Etcheverry & Zieher 1980), which causes toxin build-up in cells leading to destruction of noradrenergic terminals, and also irreversibly blocks noradrenergic transporters (Choudhary et al 2018, Ross & Stenfors 2015). To test for gross motor problems, we first examined mice in an open field arena. Wildtype mice (n=12) first underwent habituation sessions in the arena, followed by trials where both path length and velocity were measured. Animals then received either an intraperitoneal injection of 5 mg/ml DSP-4 (final concentration 50 mg/kg) (n=8) or vehicle (saline; n=4) as a control 7 days prior to further behavioural testing (Figure 3A).

**Figure 3:**
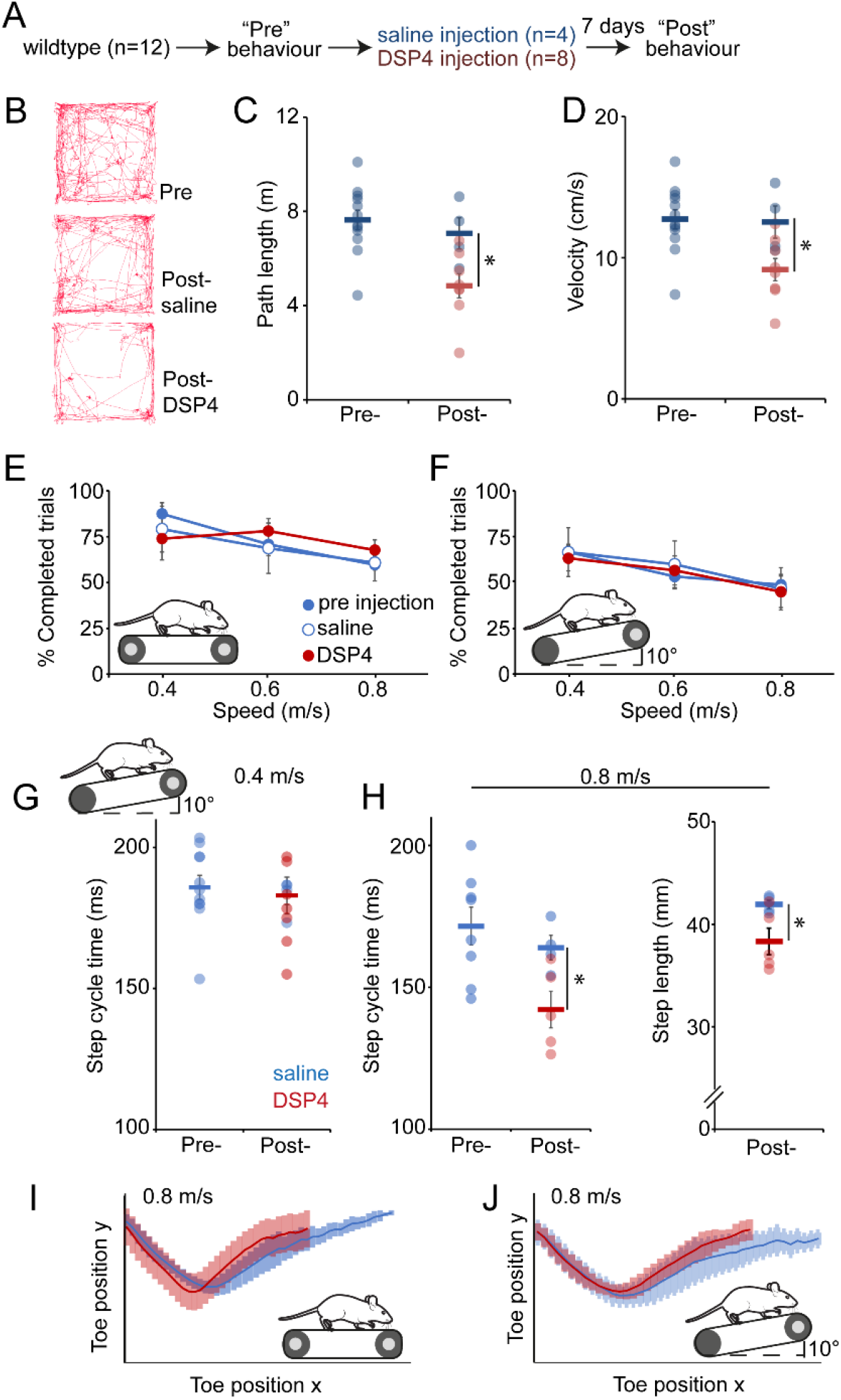
Disruption of the locus coeruleus specifically affects strenuous locomotion. (A) Experimental plan for disruption of noradrenergic signalling via the injection of the selective noradrenergic neurotoxin DSP-4. (B) Representative path lengths in 10 mins open field. (C) Overall path lengths in open field. (D) Locomotor velocity in open field. (E) Ability of control and DSP-4 injected animals to maintain consistent speed on a horizontal treadmill at 0.4, 0.6 and 0.8 metres per second. (F) Ability of wildtype and DSP-4 injected animals to maintain consistent speed on an inclined treadmill at 0.4, 0.6 and 0.8 metres per second. (G) Step cycle time on incline treadmill at 0.4 m/s. (H) Step cycle time (left) and step length (right) on incline treadmill at 0.8 m/s. (I) Toe position in xy coordinates over the step cycle at 0.8 m/s on horizontal treadmill. (J) Toe position in xy coordinates over the step cycle at 0.8 m/s on inclined treadmill. *p<0.05

Consistent with previous reports (Jones & Hess 2003) we observed a ∼25% reduction in path length when comparing the same group of animals pre- and post-toxin injection (p=0.031) and a ∼32% reduction in path length when comparing DSP-4 and saline injected animals (p=0.027; Figure 3B,C). A reduction of path length over the 5-minute open field recording could be the result of DSP-4 treated animals having overall slower locomotion, or spending more time inactive and not moving. The latter could be an indication of anxiety due to disrupted noradrenergic signalling (Brunello et al 2003). To address this, we examined the velocity of animals during periods of movement in the open field and found that DSP-4 treated animals had significantly slower movements when compared within group pre- and post-toxin injection (p=0.032) or between groups after DSP-4 or saline injection (p=0.034; Figure 3D). In addition, we did not observe any alterations in time spent in the centre of the open field arena, an indicator of anxiety levels (Supplementary Figure 3).

Next, we examined treadmill running in animals treated with DSP-4. First, we examined the ability of DSP-4 treated and control animals to maintain a constant speed on a horizontal or inclined treadmill. All groups were equally able to maintain walking on a horizontal (Figure 3E) or inclined (Figure 3F) treadmill for 3 seconds at 0.4, and 0.8 m/s. This suggests that overall locomotor ability is not dependent on noradrenergic signalling.

In addition to their ability to maintain a constant speed, we also examined aspects of locomotor kinematics across the various conditions. On an inclined treadmill at 0.4 m/s the step cycle duration of control and DSP-4 treated animals was not different between groups (pre-injection = 185.7+/-3.1 ms; DSP-4 = 183.2+/-4.2 ms; p=0.6; Figure 3G). However, at the faster speed of 0.8 m/s we noted that DSP-4 treated animals had an approximately 15-20% shorter step cycle time when compared to control groups (Figure 3H) (pre-injection = 171.5+/-6.7 ms; saline injection = 164.0+/-4.3 ms; DSP-4 injected = 142.2+/-6.4 ms; main effect of group, F=5.3, p=0.019). Similarly, DSP-4 injected animals took significantly shorter steps than saline injected controls (Figure 3H; saline= 41.9+/-0.4 mm; DSP-4= 38.3+/-1.3; p=0.049). Kinematic analysis of both horizontal and incline treadmill walking at 0.8 m/s also showed shorter steps under both conditions (Figure 3I,J).

Overall, these locomotor results show that noradrenergic signalling is important only for high intensity locomotion, namely incline running at high speed. This could indicate a role for the locus coeruleus in maintaining a high gain of locomotor activity.

### The locus coeruleus sets the level of motor response to a perturbation

We next examined whether noradrenergic signalling played a role in the generation of postural reflexes. We first looked at noradrenergic input to the LVN by examining tyrosine hydroxylase (TH; which is present in noradrenergic and dopaminergic neurons) immunostaining in the LVN. We observed extensive TH labelling in axons throughout the LVN, in close apposition to LVST-neurons labelled via cholera toxin beta (CTb) injected into the lumbar spinal cord (Figure 4A), suggesting noradrenergic input to the LVN as has been shown previously in rats (Schuerger & Balaban 1993).

**Figure 4:**
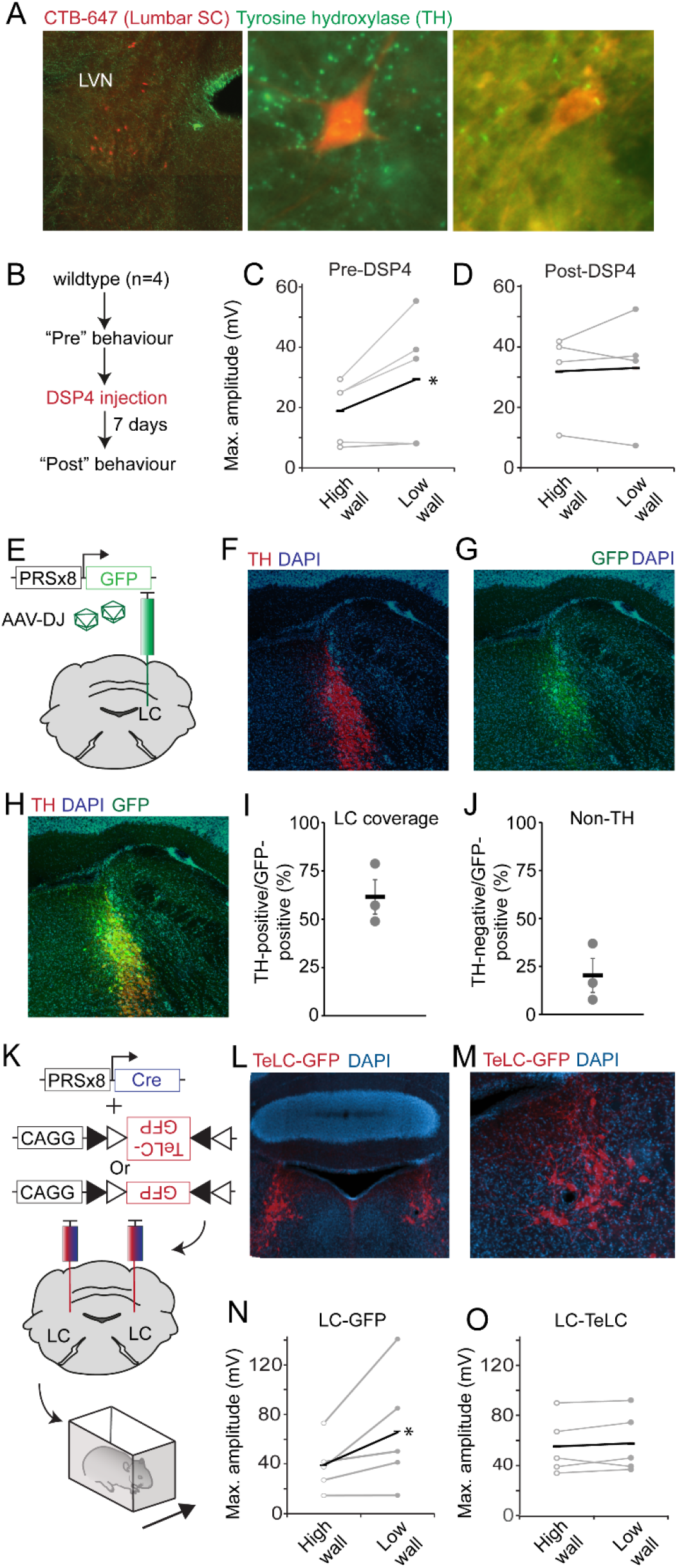
The locus coeruleus is involved in processing information regarding environmental context. (A) LVST-neurons labelled via CTB-647 injection into the lumbar spinal cord and sections stained with anti-tyrosine hydroxylase. Higher magnification images (middle and right) show apposition of tyrosine hydroxylase positive enlargements in the vicinity of LVST-neurons. (B) Experimental procedure to test effect of noradrenergic neurotoxin DSP4 on responses to lateral perturbations. (C) Comparison of peak EMG amplitudes in TA after perturbation in control animals in high and low walled conditions. (D) Comparison of peak EMG amplitudes in TA muscle after perturbation in animals following DSP4 injection in high and low walled conditions. (E) Experimental procedure for selective labelling of noradrenergic neurons in the locus coeruleus using the PRSx8 promoter. (F) Tyrosine hydroxylase immunostaining at site of injection of AAV described in (E). (G) GFP labelling and (H) merge of same site shown in (F). (I) Proportion of TH-positive neurons in the locus coeruleus that express GFP following AAV injection. (J) Proportion of TH-negative neurons expressing GFP. (K) Experimental strategy for blocking synaptic transmission from neurons in the locus coeruleus. (L) Bilateral targeting of TeLC-GFP to the locus coeruleus. (M) Higher magnification of image in (L). (N) Peak EMG responses of TA muscle in response to perturbations in high or low walled conditions in animals expressing GFP in the locus coeruleus. (O) Peak EMG responses of TA muscle in response to perturbations in high or low walled conditions in animals expressing GFP-TeLC in the locus coeruleus.

Next, we examined whether pharmacological disruption of noradrenergic neurons would affect the EMG response to a perturbation in different environmental conditions. Again, we systemically administered DSP4 to disrupt noradrenergic neurotransmission (Figure 4B). In the TA muscle, before administration of DSP4, EMG peak amplitudes were greater in low walled (34.3±13.9 mV) than high walled (16.0±4.2 mV; p=0.044) conditions. After systemic administration of DSP4, similar peak EMG amplitudes were recorded in mouse hind limb muscles in the high (32.2±6.4 mV) and low (35.3±14.6 mV) walled conditions following perturbation (p=0.73; Figure 4B-D). These results are consistent with a potential role for noradrenergic signalling in setting the gain of a postural response.

Although DSP-4 is a potent neurotoxin for noradrenergic neurons of the locus coeruleus resulting in ∼90% neuron ablation, it has been reported to ablate up to 50% of neurons in other noradrenergic nuclei, including those which innervate the spinal cord (Lyons et al 1989). To ascertain whether the alterations in postural corrections are indeed coordinated by the locus coeruleus, and not other noradrenergic nuclei, we used a viral strategy for selective blockade of neurotransmission from only those neurons. We took advantage of a previously reported noradrenergic selective promoter PRSx8, a synthetic promotor containing Phox2B binding motifs (Hwang et al 2001), which we cloned and packaged into an AAV. We first assessed the specificity of the PRSx8 promoter in an AAV by placing it upstream of a GFP reporter which was packaged into an AAV-DJ capsid (Figure 4E). Stereotaxic injection of this vector into the LC of wildtype mice resulted in 61.6%+/-8.9% of LC neurons expressing GFP (Figure 4F-I). In contrast, 20.3+/-8.6% of GFP-positive neurons did not express tyrosine hydroxylase (though this could represent neurons expressing low levels of tyrosine hydroxylase) (Figure 4J).

We next packaged an AAV expressing cre recombinase under control of the PRSx8 promoter and co-injected this along with an AAV containing a cre-conditional tetanus toxin light chain fused to GFP, or GFP alone (Figure 4K)(Murray et al 2011). Tetanus toxin light chain prevents synaptic transmission through disruption of vesicle docking. In these experimental conditions synaptic transmission is selectively blocked from noradrenergic neurons in the locus coeruleus only (Figure 4K,L,M). Control animals expressing GFP in the locus coeruleus showed an increased EMG amplitude in response to perturbation in low walled vs. high walled conditions in the TA muscle (high mean max amplitude 38.8±4.4mV; low mean max amplitude 66.5±9.7; n=5 mice; p=0.048 Figure 4N). However, blockade of synaptic transmission resulted in highly similar EMG responses in high and low walled conditions (high mean max. amplitude 53.0±4.7mV; low mean max. amplitude 53.6±5.3; n=5 mice; p=0.89 Figure 4O). Therefore, in agreement with our data using DSP-4, selective blockade of synaptic transmission from the locus coeruleus results in postural responses no longer being tuned to the environment. Overall, these results support a model whereby fast postural corrections are initiated by the LVN and the locus coeruleus sets the level of this response according to the environmental context.

## Discussion

In order to maintain upright posture, the nervous system must quickly respond to mechanical perturbations that impact the body and produce a counteracting motor output. The motor response can change according to the environmental context and can be heavily influenced by perceived threat. Here we have shown that the locus coeruleus can alter the gain of vestibulospinal reflexes that maintain balance following a postural perturbation. This noradrenergic influence on motor circuits provides a potential link between forebrain regions that interpret environmental context, and the sensory-motor reflex pathways of the brainstem and spinal cord.

In response to an unexpected postural challenge, or perturbation, animals and humans must generate a motor response that counteracts the displacement. Rather than this being a simple motor reflex, it is well known that these postural corrections are adapted to suit a range of environmental and spatial contexts (Bolton 2015, Jacobs & Horak 2007, McCollum et al 1996). For example, the gain of postural reflexes can be altered in humans when they are standing on the edge of a high platform (Horslen et al 2014). Evolutionarily, this mechanism could indicate the nervous system focussing more on postural reflexes under circumstances where the cost of falling is extremely high, or to maintain a stable body position under threatening conditions (Balaban 2002, Horslen et al 2018). To probe the circuit mechanisms that link the environment to motor reflexes we developed a mouse behavioural assay that allows for adaptation of postural reflexes in different environmental contexts. When in unfamiliar environments mice tend to prefer enclosed spaces, as observed by thigmotaxis in open field arenas or the preference for enclosed areas on a plus maze (Kulesskaya & Voikar 2014). We reasoned that mice in more exposed environments would experience higher threat and stress levels than when in a more enclosed spaces, and that this threat could mimic that experienced by humans when standing at height (Horslen et al 2014). Indeed, by altering the height of the surrounding walls on a moving platform we could alter the motor correction and muscle response. Specifically, when in a more open environment the amplitude of the muscle response to the perturbation (as measured via EMG) was increased, resulting in a greater postural correction and reduced body sway even when the perturbation was of equal magnitude across the two environments. Using selective circuit manipulations we showed that this effect is mediating by noradrenergic signalling from the locus coeruleus.

Noradrenaline is known to be released in response to stress or in situations of heightened arousal (Ross & Van Bockstaele 2020) and circuits linking the locus coeruleus and the forebrain have indicated that noradrenaline has a role in attention and vigilance (Sara 2009). However, this is not the only role of noradrenaline in in the nervous system. Classical studies in decerebrate animals have indicated that noradrenaline can play a facilitatory role in postural responses (Pompeiano 2001), can influence the gain of vestibulospinal reflexes (Pompeiano et al 1991) and cessation of activity in the locus coeruleus has been observed during loss of muscle tone in cataplexy (Wu et al 1999). Noradrenergic signalling can be facilitatory or inhibitory, depending on the postsynaptic receptor complement. In the cat microiontophoretic injection of noradrenaline into the lateral vestibular nucleus increases resting discharge of the neurons, whereas conversely in the adjacent medial vestibular nucleus it decreases resting activity (Kirsten & Sharma 1976). The locus coeruleus can also influence spinal circuits directly via coeruleospinal pathways, though this pathway has been predominantly implicated in nociceptive processing (Suarez-Pereira et al 2022). In addition, the locus coeruleus itself is known to receive input from a diverse array of brain regions including prefrontal cortex, hypothalamus and amygdala (Schwarz et al 2015) which could inform the postural control system of the surrounding environment. These sensory-motor relationships provide a potential circuit whereby higher order information regarding the nature of the surrounding environment is routed through the locus coeruleus to motor pathways of the LVN.

Disruption of noradrenergic signalling had minor effects on locomotor activity. In agreement with previous studies we observed a reduced open field path length after injection of DSP-4 (Harro et al 2000). Additionally, in a task that required increased locomotor effort we did observe that mice with disrupted noradrenergic signalling had altered stepping (shorter and faster steps) when compared to control animals. Potentially, this could indicate noradrenergic signalling also feeds into locomotor circuits under circumstances that require an enhanced motor output.

Clinically, degeneration of the locus coeruleus in Parkinson’s disease has been linked to problems with balance (Gesi et al 2000, Grimbergen et al 2009). Some aspects of Parkinsonian postural responses – such as an increased amplitude of the medium latency postural response – are reminiscent of the results obtained in our study (Rinalduzzi & Curra 2015, Rinalduzzi et al 2015). Additionally, Parkinsonian patients show a reduction in locomotor step length and velocity (Chastan et al 2008). Furthermore, drugs which raise noradrenaline levels are related to increased risk of falls (Park et al 2015). Interestingly, gait disorders and an increase in falls are particularly clinically challenging because they do not tend to improve with the common therapeutic interventions levodopa and deep brain stimulation (Devos et al 2010a, Devos et al 2010b). However, treatment of a rat model of Parkinson’s disease with noradrenaline reuptake inhibitors alone or in combination with alpha2 receptor antagonists can rescue some motor deficits (Yssel et al 2018) of the disorder. Coupled with our results, this could indicate that disruption of noradrenergic signalling in Parkinson’s disease could contribute to the balance and motor deficits by altering the activity of vestibulospinal pathways.

At a circuit level, we suggest two potential mechanisms whereby locus coeruleus activity could influence postural control. First, in the event of an unexpected perturbation, the locus coeruleus uses information about the environment to fire phasically with the LVN, which then implements the motor correction at the level of the spinal cord. In an alternative model, the locus coeruleus is informed of the change in context prior to any perturbation and alters motor output either via the LVN or directly through spinal motor neurons via the coeruleospinal pathway. This second model would be consistent with the known role of the locus coeruleus in stress, where sensitivity to hormonal changes means that LC firing tends to increase tonically rather than in response to discrete stimuli (McCall et al 2015, Poe et al 2020, Valentino & Van Bockstaele 2008), and so it may have a similar role in the postural control system.

## Methods

### Animals

All experiments were performed under UK Home Office license according to the United Kingdom Animals (Scientific Procedures) Act 1986. Both male and female C57BL/6J mice 12-20 weeks old were used.

### Surgical procedures

#### EMG implantations

Bipolar EMG electrodes were fabricated and implanted into the hindlimb muscles gastrocnemius, tibialis anterior, semitendinosus and vastus laterals using techniques described previously (Akay et al 2014). Briefly, mice were anesthetised in isoflurane (4% induction; 0.5-2% maintenance). The neck and hindlimb were shaved with incisions made at the neck and directly above the muscles to be implanted. Custom made bipolar electrodes were passed under the skin from the neck to the hindlimb and implanted into the appropriate muscles. Animals were allowed to recover for at least three days before beginning behavioural experiments. Prior to beginning perturbation experiments animals walked on a treadmill at a constant speed of 0.3 m/s while EMG was recorded to ensure the EMG signals were of sufficient quality and the expected flexor-extensor alternation pattern was obtained.

#### Stereotaxic injections

Stereotaxic injections into the LVN were carried out as described previously (Murray et al 2015). Briefly, mice were anesthetised with isoflurane (4% induction; 1-2% maintenance). An incision was made in the skin above the scalp and bregma and lambda visualised. A small burr hole was made above the injection site and AAV was injected using a Nanoject II or Nanoject III (Drummond) and a pulled glass pipette. Stereotaxic coordinates relative to bregma were as follows: lateral vestibular nucleus, anterior/posterior -6.05 mm; lateral 1.38 mm; depth from brain surface -4.5, -4.4 and -4.3 mm with each depth receiving 100 nl of AAV. Locus coeruleus, anterior/posterior -5.4 mm; lateral 0.85 mm; depth -3.85, -3.75, 3.65 mm with each depth receiving 50 nl of AAV. Following AAV injection the skin was closed with Vicryl Rapide sutures. Behavioural experiments commenced no sooner than 14 days after AAV injection to allow sufficient time for the transgene to be expressed.

### Behavioural Procedures

#### Lateral perturbations

Mice were anaesthetised using isoflurane to attach recording connectors then placed in enclosures 10 cm by 6 cm with walls of either 3.5 cm or 7 cm in height. The enclosure was situated atop a treadmill such that it could move laterally as the treadmill belt rotated. A TTL pulse was used to induce a reproducible movement of the treadmill which caused a lateral displacement of the enclosure of 115 mm with a peak acceleration between 0.53 and 0.54 ms, reached 50 ms from movement onset, and the entire movement lasted 140 ms. The order of testing each mouse in the different wall heights was randomised to control for any habituation to the perturbation. Trials were monitored using an overhead camera (see recording techniques below) and only trials where the animal maintained a consistent starting position were analysed. Partial obstruction of the overhead view of the animals by EMG wires and fiber optic cables prevented precise tracking of body displacement in the same animals that had EMG or fibre optic implants.

#### Treadmill locomotion

Locomotion was tracked on a custom-built treadmill (Electronics workshop, Zoological Institute, University of Cologne). Video recording of the right-side view of the mouse was carried out with a high speed camera (Ximea) recording at 200 frames per second. For incline running experiments the front of the treadmill apparatus was raised such that the treadmill belt had an upward gradient of 10 degrees.

#### Open field

Mice were allowed to freely explore a square white Perspex arena for a period of 10 minutes while being recorded by an overhead video camera recording at 30 frames per second. To avoid effects of habituation on experiments mice were exposed to the arena for 10 minutes on the three days prior to experimental recordings.

### Recording techniques

Muscle activity was amplified using custom built pre-amplifiers and amplifier (University of Cologne), digitised using a 1401-3 (Cambridge Electronic Design), and recorded using Spike2 (Cambridge Electronic Design). Video was captured using an overhead Logitech Brio Webcam and side mounted Doric Behaviour Tracking camera. Reflective markers were attached to the enclosure and the mouse’s head to track relative head position during perturbation. In some trials animals were also recorded in the arena for ∼10 seconds before and after perturbations in order to provide data for respiration rate.

### Data Analysis

Video was analysed using MaxTraq Software (Innovision Systems). For treadmill running the toe and ankle from the right hindlimb were tracked while the mouse matched the speed of the treadmill. To provide an internal measure of limb movement relative to the body the eye and tip of the ear were also tracked. Tracking data was exported to Microsoft Excel for plotting and further analysis. Conditions pre and post drugs were compared using student’s paired T-test.

To measure head displacement during perturbation experiments video data was again analysed using MaxTraq Software (Innovision Systems). A fixed point on the top of the animal’s head was tracked along with a fixed point on the arena. To measure head displacement the angle between the head and arena points was measured from the onset of the perturbation through to ∼500 ms after the perturbation when the head had generally returned to its starting position.

EMG signals were recorded in Spike2, rectified and DC variation removed. Maximum amplitude responses within 100ms of perturbation were exported to Microsoft Excel. Student’s t tests were used for comparison of EMG signals and for quantification of ablation efficiency. P < 0.05 was considered significant.

For analysis of respiration rate breaths using high speed video taken before or after were counted by an experimenter naïve to the experimental condition.

### Viruses

For AAV constructs using the PRSx8 promoter, the PRSx8 promoter sequence based on Hwang et al., 2001 was de novo synthesised (Life Technologies) and inserted into the pAM AAV genome vector (Murray et al., 2011) upstream from the transgene via KpnI and BamHI restriction sites. The sequence of the synthesised PRSx8 promoter was as follows:

5’-AGCTTCCGCTAGACAAATGTGATTACCCCCGCTAGACAAATGTGATTACCCGCGCTAGACAAATGTGATTACCCCGCTAGACAAATGTGATTACCCCCCGCTAGACAAATGTGATTACCCCCGCTAGACAAATGTGATTACCCGCGCTAGACAAATGTGATTACCCCGCTAGACAAATGTGATTACCCCCGACCAGGGCATAAATGGCCAGGTGGGACCAGAGAGCTCACCCCAGCCGACTCTAG

-3’

Optogenetic inhibition was achieved with an AAV (DJ serotype) expressing archaerhodopsin (ArchT) (Han et al 2011) obtained via Addgene (plasmid #29778).

AAVs were packaged via calcium phosphate transfection of HEK293 cells as detailed in (McClure et al 2011) and purified using an AAVpro purification kit (Takara Bio).

### Drugs

DSP4 blocks noradrenaline transporters, which reduces levels of noradrenaline and eventually kills noradrenergic neurons. DSP4 (Sigma C8417) was dissolved in sterile saline and injected intraperitoneally to produce a final concentration of 50 mg/kg. First injection was carried out immediately after recording control behavior with a second dose after 4 days. Behavioural testing was carried out 2-4 days after administration to ensure maximal decrease in noradrenergic neurons (Choudhary et al 2018).

### Histology

After recording, mice were transcardialy perfused using ice cold 4% paraformaldehyde (PFA). Brains were harvested and fixed in 4% PFA overnight before being transferred to phosphate buffer solution. 50 µm thick coronal brain sections were cut on a vibratome (Leica) and mounted on Superfrost+ glass slides for histological processing. Antibodies used were as follows: Rabbit to tyrosine hydroxylase (1:500) (Abcam, catalogue number AB152); Goat anti GFP (1:1000) (Abcam catalogue number ab167453); Rabbit anti aldolase C (1:500) (Life Technologies, catalogue number PA551883).

## Acknowledgements

We thank Dr. Turgay Akay (Dalhousie University), Dr. Niccolo Zampieri (MDC Berlin), Prof. Rob Brownstone (University College London) and members of the Murray lab for comments on the manuscript. This work was supported by Wellcome grant 219627/Z/19/Z, ‘Sainsbury Wellcome Centre Core Grant” and Gatsby Charitable Foundation grant GAT3755 “Sainsbury Wellcome Centre (SWC) core grant”. An image from SciDraw made by Gil Costa (scidraw.io) was used in Supplementary Figure 1B.

