## Supplementary Material for "The locus coeruleus directs sensory-motor reflex amplitude across environmental contexts"

**Witts et al.**

### Supplementary Figures

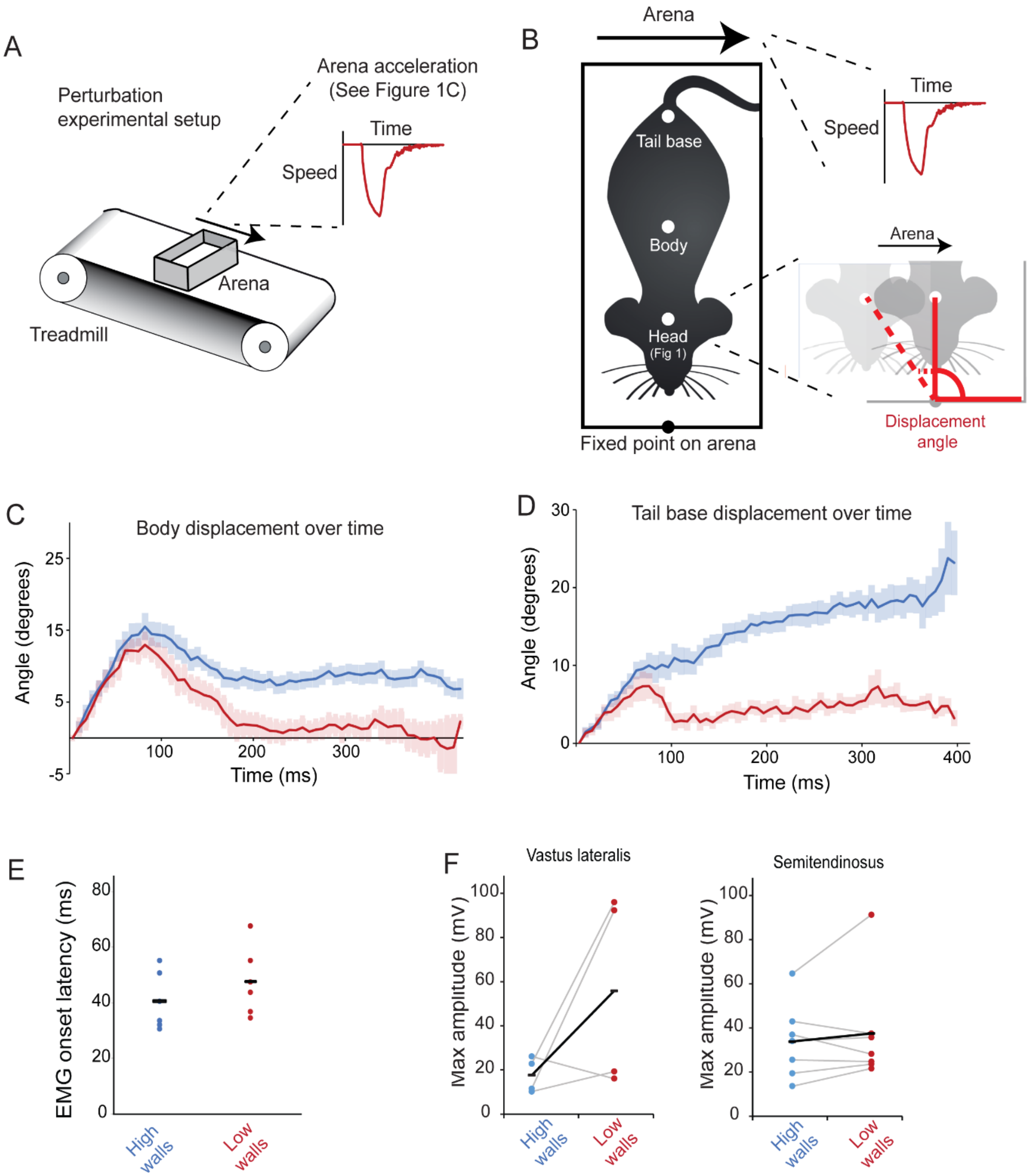


**Supplementary Figure 1:** (A) Schematic showing experimental setup and perturbation acceleration. (B) Schematic showing placement of tracking dots and measurement of displacement angle. (C) Body displacement angle following perturbation. (D) Tail base displacement following perturbation. (E) Onset latency of muscle response following perturbation in high and low wall conditions. (F) Peak EMG amplitude of VL and ST muscles during perturbation in high and low walled conditions. Each point is the mean of trials from individual experimental animals with the overall mean represented by black lines.

A HIGH WALLS

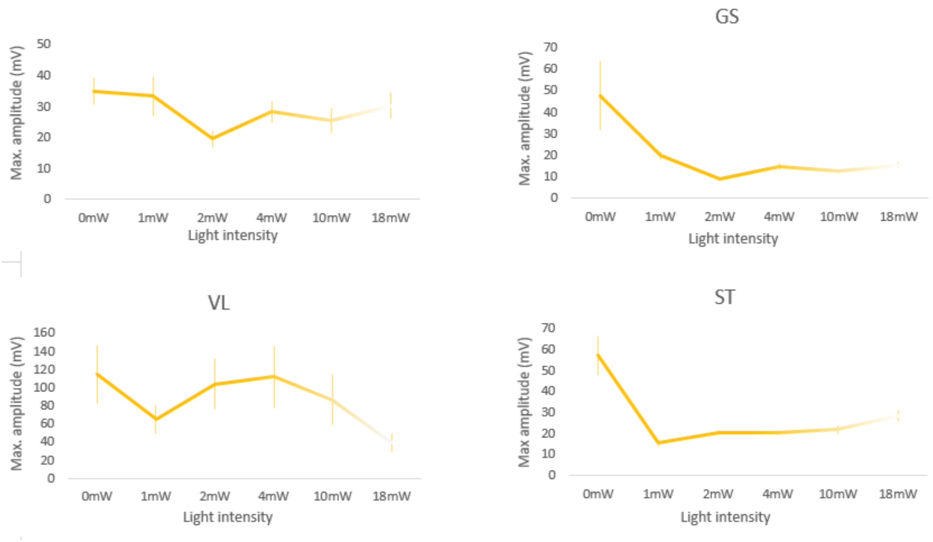


B LOW WALLS

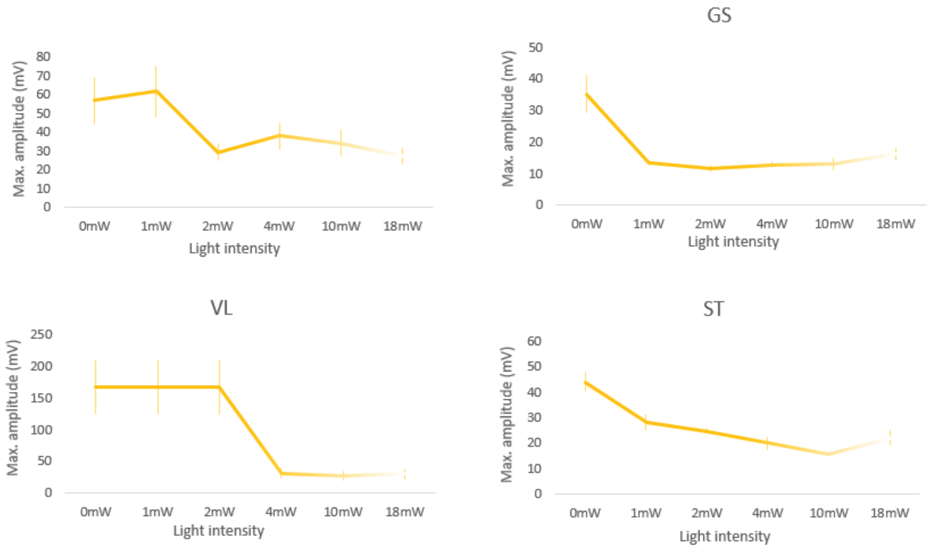


**Supplementary Figure 2:** (A) Mean peak EMG amplitude in high wall condition in four muscles after perturbation with inhibition of LVN neurons at different light intensities. (B) Mean peak EMG amplitude in low wall condition in four muscles after perturbation with inhibition of LVN neurons at different light intensities. (n= 4 animals, each light intensity repeated twice with order of light intensity varied between trials).

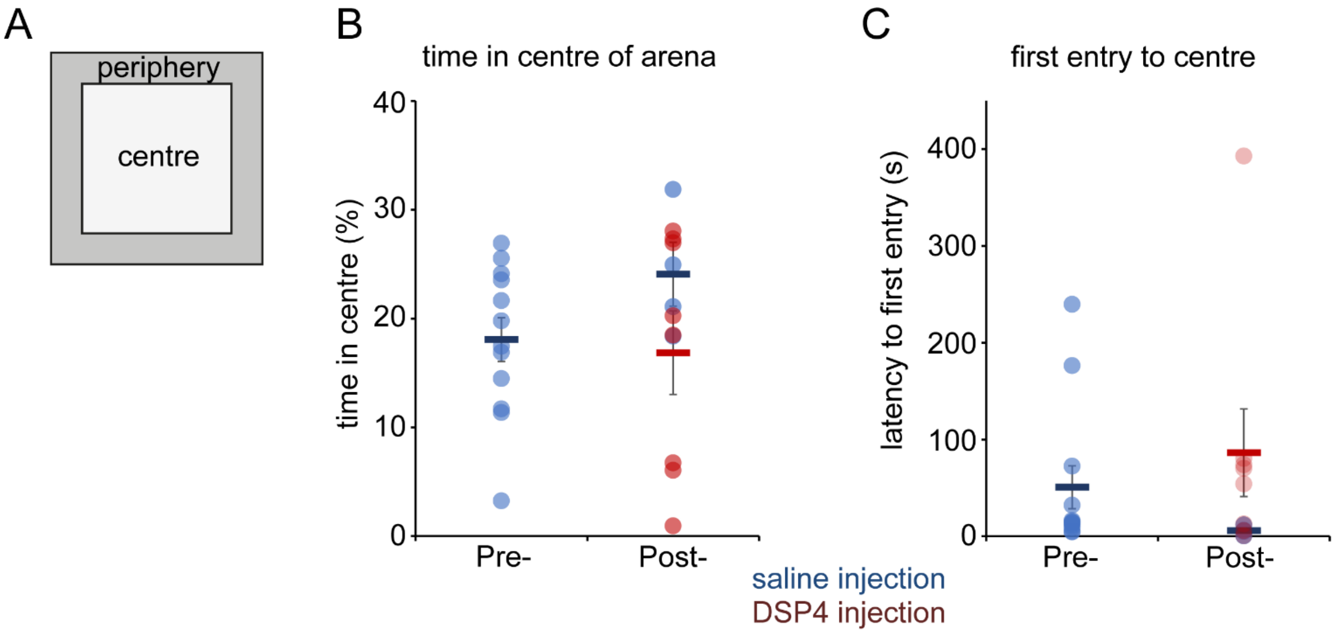


**Supplementary Figure 3:** (A) Schematic of open field arena. (B) Percentage of time spent in centre of arena pre- and post- DSP4 or saline injection. (C) Latency of first entry to centre of arena pre- and post- DSP4 or saline injection.
